# Acute myocardial injury in *mdx* hearts ameliorated by ARB but not ACE inhibitor treatment

**DOI:** 10.1101/765602

**Authors:** Tatyana A. Meyers, Jackie A. Heitzman, DeWayne Townsend

## Abstract

Duchenne muscular dystrophy (DMD) is a devastating muscle disease that afflicts males due to the loss of the protein dystrophin, resulting in muscle deterioration and cardiomyopathy. Dystrophin’s absence causes increased membrane fragility, myocyte death, and tissue remodeling. Inhibition of angiotensin signaling with ACE inhibitors or angiotensin receptor blockers (ARBs) is a mainstay of DMD therapy, with clinical guidelines recommending starting one of these therapies by the age of 10 to address cardiomyopathy.

Using the *mdx* mouse model of DMD, we previously showed that isoproterenol causes extensive damage in dystrophic hearts, and treatment with the ARB losartan starting only 1 hour before isoproterenol dramatically reduced this myocardial injury. In the present study, we probed whether ACE inhibitors, which are more frequently prescribed, can deliver similar protection. Surprisingly, lisinopril treatment initiated 1 hour before isoproterenol failed to demonstrate any effect on injury in *mdx* hearts. Further, with a 2-week pretreatment, only losartan significantly lowered *mdx* cardiac injury, without any benefit associated with lisinopril treatment. These results confirm the ability of ARBs, but not ACE inhibitors, to prevent acute injury in mouse hearts, and prompt the question whether ARBs should be more frequently used for DMD cardiomyopathy because of these potential protective actions.

## 1. Introduction

Duchenne muscular dystrophy (DMD) is a devastating muscle wasting disease that results in the loss of skeletal muscle function and development of cardiomyopathy. This X-linked disease occurs almost exclusively in males at a rate of roughly 1:5000 live male births [1,2]. The absence of the protein dystrophin leads to increased sarcolemmal fragility and dysregulation of signaling processes associated with proper dystrophin localization, triggering myocyte necrosis and replacement with fibrosis. DMD is lethal in young adulthood, with death typically occurring from respiratory or heart failure. In recent decades, advancements in symptomatic respiratory therapies have resulted in prolonged patient lifespans, leading to higher rates of advanced heart disease among individuals with DMD [3].

The cardiomyopathy of DMD is a progressive disorder, with the earliest signs of disease processes appearing before the age of 10, and extensive myocardial fibrosis and contractile deficits marking the end stage of the disease [2,4–6]. Attempts to slow the progression of heart disease in patients with DMD include the recommendation that chronic treatment with ACE inhibitors (ACEIs) or angiotensin receptors blockers (ARBs) be initiated by 10 years of age or earlier, with the goal of mitigating fibrosis and adverse remodeling that follows cardiomyocyte loss in the dystrophic heart [7,8]. Both drugs aim to limit the deleterious processes associated with angiotensin II (AngII) signaling in the heart and muscle, which include increased production of reactive oxygen species (ROS), profibrotic gene transcription, and myocyte death [9–12]. ACEIs interfere in these processes upstream of AngII, hindering its production by blocking the ability of angiotensin-converting enzyme (ACE) to cleave angiotensin I into AngII. Conversely, ARBs limit the effects of AngII after it is formed by blocking activation of angiotensin II type 1 receptor (AT_1_R), a key mediator of many of AngII’s harmful effects. Several early clinical trials demonstrated improved cardiac function in DMD patients using ACEIs, paving the way for ACEIs being viewed as the top candidate for heart failure therapy [13,14]. Additional trials went on to show that ARBs were just as effective as ACEIs when used in patients whose cardiac health has begun to decline, and that ARBs are better tolerated by patients, but ACEIs continue to be used more widely in clinical management of DMD [15–17].

Recently, our group used the *mdx* mouse model of DMD to demonstrate that a single bolus injection of the β-adrenergic receptor agonist isoproterenol (Iso) results in acute damage to 18.9±0.3% of the *mdx* myocardium, relative to 6.3±2% injury in wild type mice [18]. This experimental model of increased cardiac workload aims to approximate the bouts of stress experienced by DMD patient hearts during surgical procedures [19], with the subsequent injury likely resulting from the exacerbation of the ongoing disease mechanisms of the dystrophic heart [20,21]. Importantly, we reported that dystrophic *mdx* mice acutely treated with the ARB losartan shortly before this injurious stimulus show a striking degree of protection from myocardial injury. High-dose losartan treatment initiated only 1 hour prior to the bolus of Iso resulted in a 2.8-fold reduction in *mdx* myocardial injury, seemingly preventing the accumulation of myocyte damage in the dystrophic heart [18]. Interestingly, acute losartan did not have a significant effect on Iso-induced damage in wild type hearts, resulting in similarly low injury area in *mdx* and healthy hearts (6.8±2% vs. 4.9±1%, respectively) [18]. This demonstrated that dystrophic hearts are uniquely vulnerable to AT_1_R-based exacerbation of myocardial injury via a mechanism that plays a negligible role in healthy hearts.

This discovery of losartan’s ability to prevent dystrophic heart injury has prompted the question whether ACEIs would exert a similar effect. While it is well-documented that both classes of drugs are effective at limiting the fibrosis and remodeling that follow injury, the preservation of functional dystrophic myocardium is of utmost importance in the management of cardiac disease in patients with DMD. To determine the effectiveness of ACEIs in preserving dystrophic myocardium, here we present our previously published Iso-induced injury protocol with the ACEI lisinopril. Our new findings demonstrate that neither acute nor chronic treatment with lisinopril shows the protective capacity to limit Iso-induced myocardial loss in dystrophic hearts that is present with ARB treatment.

## 2. Methods

### 2.1. Animals

The wild type strain *C57BL/10SnJ* (C10) and the dystrophic strain *C57BL/10ScSn-Dmd*^*mdx*^*/J* (*mdx*) were bred and maintained at the University of Minnesota from breeding stock obtained from Jackson Laboratories. All mice were 4-6 mo of age at the time of experiments. Since DMD almost exclusively affects males, male mice were used in these studies. All animal procedures were approved by the University of Minnesota Institutional Animal Care and Use Committee, and were performed in compliance with all relevant laws and regulations.

### 2.2. Isoproterenol studies

(-)-Isoproterenol hydrochloride (Iso; Sigma #I6504) was dissolved in saline to a concentration of 6 mg/ml and sterile-filtered prior to injection. The sterile Iso solution was stored protected from light at 4°C for up to 3 days, and discarded at the first sign discoloration indicative of degradation. Mice received a single intraperitoneal bolus injection of 10 mg/kg Iso in a volume of 40-70 µl, and hearts were harvested 30 hours after the Iso injection, as previously described [18].

### 2.3. Acute lisinopril study

A subset of mice received acute lisinopril treatment starting 1 hour prior to Iso and continuing after Iso, terminating at the 30-hour timepoint. Lisinopril powder (Cayman Chemical #16833) was dissolved in sterile saline on the evening prior to Iso injections to a concentration of 12 mg/ml, and stored protected from light at 4°C. At the start of the light cycle (7 am) on the day of the study, the mice received a 20 mg/kg loading dose of lisinopril via IP injection in a volume of 40-70 µl, followed by a 10mg/kg Iso injection 1 hour later. At the end of the light cycle (7 pm), the mice received another bolus of 20 mg/kg lisinopril to span the dark cycle. At the end of the dark cycle, they received a final dose of 20 mg/kg lisinopril, and hearts were harvested 7 hours later (30 hours after the Iso bolus). This dosing regimen was based on the serum half-life of lisinopril, and designed to minimize troughs in drug serum levels. Littermate *mdx* and wild type mice were distributed evenly between the groups receiving Iso with lisinopril or Iso only.

### 2.4. Lisinopril and losartan pretreatment studies

To account for the possibility that additional protective effects may be revealed with pretreatment, an additional subset of mice was assigned to drink either lisinopril or losartan dissolved in water for 2 weeks prior to Iso-induced injury. Lisinopril was given at a concentration of 264 mg/L in 1% dextrose water with an average consumed dose of 40 mg/kg daily [22]. Losartan was given at a concentration of 600 mg/L in 3% dextrose, with an average consumed dose of 80 mg/kg daily [23]. The 2-fold difference in doses of losartan and lisinopril is reflective of the similar 2- to 3-fold difference in doses of these drugs in human clinical applications. Drug-containing water was replaced 3 times per week, and each mouse was weighed before and during the pretreatment at weekly intervals [22].

The physical discomfort experienced by mice following Iso administration can disrupt their water and food intake for up to 24 hours, so the mode of drug delivery was switched from the drinking water to IP injections to ensure that therapeutic levels are maintained during that time. At the start of the light cycle on the 15^th^ day after the start of pretreatment, mice received fresh tap water and a single a single 10mg/kg Iso IP injection. For the next 30 hours, the mice pretreated with lisinopril received the same booster injections of lisinopril as described in section 2.3 above, and mice pretreated with losartan received booster injections as described previously [18]. Mice were sacrificed and hearts were harvested 30 hours after the Iso injection.

### 2.5. Histopathology

Histopathology studies were carried out as previously described [18]. Briefly, fresh excised hearts were cut in half along the transverse plane and placed into OCT medium to be frozen in liquid nitrogen-cooled isopentane. Heart sections were cut to 7 µm, and the slides were stored at −80°C before staining. All staining was performed on unfixed tissue. The following reagents were used for IF staining: goat serum for blocking (Jackson ImmunoResearch # 005-000-121, 10%), goat anti-mouse IgG (H+L) secondary antibody (Invitrogen #R37121, 1:200), WGA AlexaFluor conjugate (ThermoFisher, 5µg/ml), and ProLong Gold Antifade Mountant with DAPI (ThermoFisher). After blocking, staining was performed as a single step at room temperature for 1 hour, flanked by three 5-minute washes in PBS.

Only hearts from mice that survived the initial 24 hours were analyzed for injury area, due to the difficulty of quantifying the poorly-defined early IgG-positive injury present in hearts that succumb earlier in the time course. To ensure that this approach would not skew results, we confirmed that the mortality rates of 20-37% among *mdx* mice and 5-14% among wild type mice did not differ significantly between treatments.

### 2.6. Microscopy and image analysis

All imaging was performed in NIS Elements software on a Nikon Eclipse Ni-E upright epifluorescent microscope with motorized stage and filter wheel. Whole heart montages were collected as a stack of three fluorescent channels at a resolution of 0.92 µm/pixel. IgG-positive injury area was determined using Fiji by subtracting the WGA channel (staining extracellular matrix) from the IgG channel to better distinguish intra-myocyte IgG signal [24]. This lesion area was then quantified by thresholding the resulting IgG image for total lesion area, and normalizing it to total heart section area.

### 2.7. Patient anti-angiotensin drug information

Electronic medical records from patients diagnosed with Duchenne muscular dystrophy at the University of Minnesota were surveyed for prescriptions of either ACEIs or ARBs. In total, we examined the records of 95 DMD patients seen from 2009-2019. The 30 patient records that did not specifically mention an ACE inhibitor or an ARB were excluded from the analysis. Several patients had received both ACEIs and ARBs, and in these cases they were classified by the most recent prescribed class of medication. No patients were receiving both drug classes at the same time. Collection and analysis of these data were performed in accordance with an IRB-approved protocol.

### 2.8. Statistics

Statistical tests were performed using Prism 7 (GraphPad Software). Preclinical study statistical comparisons were made using two-way ANOVA with Sidak post-hoc test. Ages from patient data were compared using the non-parametric Mann-Whitney test. All graphs display the mean ± standard error.

## 3. Results

### 3.1. Acute treatment with lisinopril has no effect on Iso-induced cardiac injury area

Acute ACE inhibition with lisinopril initiated 1 hour prior to the bolus of Iso failed to decrease IgG+ cardiac injury area in both *mdx* hearts and wild type hearts (Fig. 1). As previously shown, *mdx* hearts stressed with a 10 mg/kg bolus of Iso displayed very high injury compared to wild type hearts (20.2±2% vs. 5.3±2%), and this difference persisted with ACE inhibition (16.2±2% vs. 6.4±2%) (Fig. 1 B and C). This result stands in stark contrast to our earlier published findings demonstrating that acute losartan treatment has a robust capacity for myocardial injury reduction in dystrophic hearts [18].

**Figure 1:**
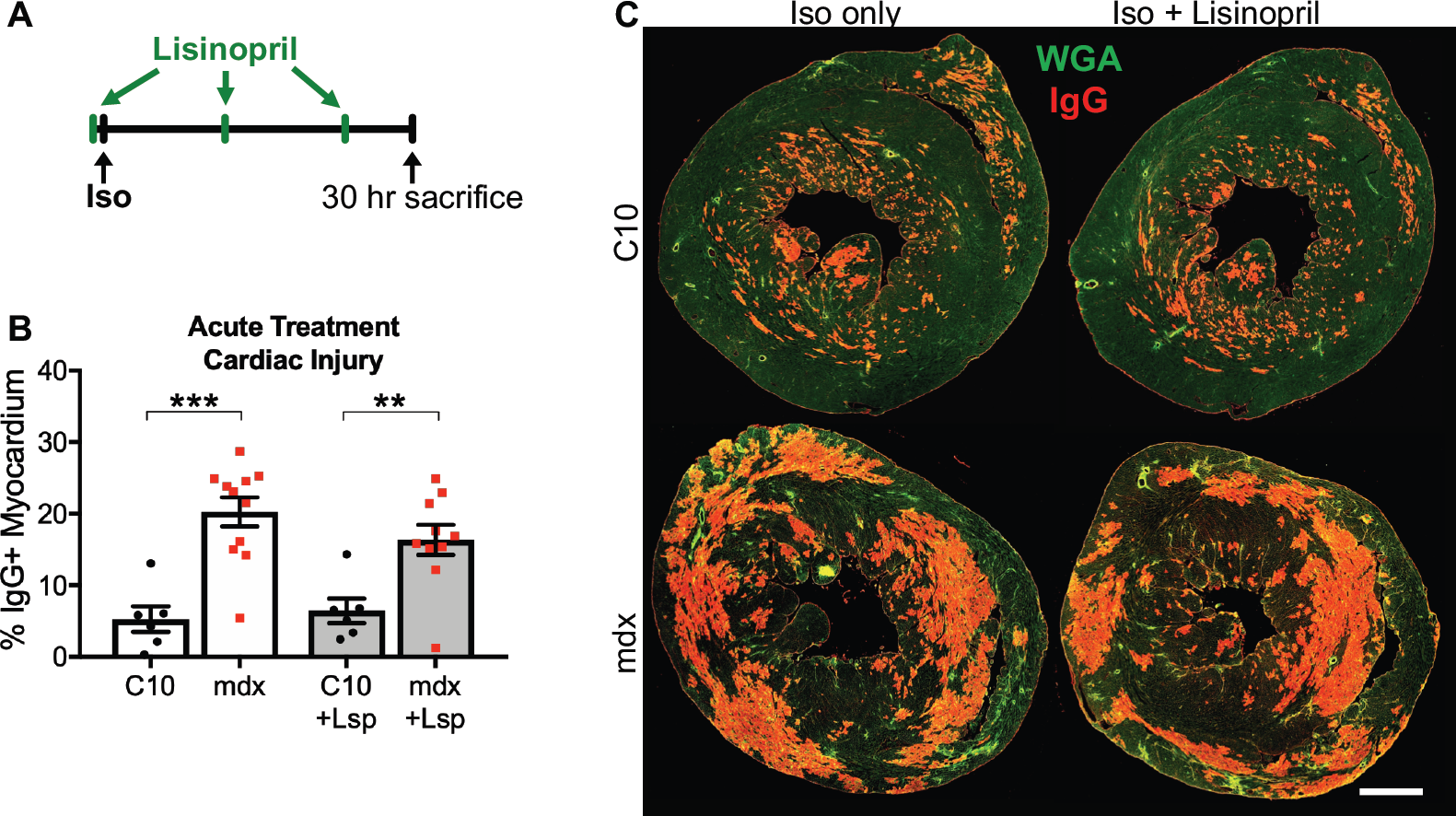
Acute ACE inhibition is insufficient to protect the heart from Iso-induced injury. **(A)** The ACE inhibitor lisinopril was given as a 20 mg/kg IP injection 1 hour prior to a 10 mg/kg bolus injection of isoproterenol (Iso). Booster injections of lisinopril were given twice at 12-hour increments to maintain drug serum levels. **(B)** Dystrophic (*mdx*) hearts displayed 3.8-fold higher myocardial injury than wild type (C10) hearts 30 hours post-Iso. Acute treatment with lisinopril (Lsp) did not significantly change Iso-induced injury area in either group (***p<0.001, **p=0.003; C10 n=6 per group; *mdx* n=10–11 per group). **(C)** Representative images of data in panel B. Whole heart transverse sections are shown with WGA (green) marking the tissue area and IgG (red) indicating areas of myocardial injury. Scale bar = 1 mm.

### 3.2. Pretreatment with losartan, but not with lisinopril, significantly reduces Iso-induced injury in *mdx* hearts

To address the possibility that lisinopril may require a period of pretreatment to exert a protective effect, *mdx* and C10 mice were placed on a 2-week course with either lisinopril or losartan delivered in their drinking water before repeating the bolus injection protocol (Fig. 2A). In agreement with our earlier work, losartan pretreatment exerted a significant protective effect in dystrophic hearts, reducing *mdx* cardiac injury to 12.2±3% without any effect in C10 hearts (5.7±1% injury area) (Fig. 2 B and C). In contrast, pretreatment with lisinopril still did not reveal any protective effect on myocardial damage in mouse hearts. *Mdx* mice pretreated with lisinopril and injected with a bolus of Iso showed 20.9±2% IgG-positive cardiac injury, and C10 mice in the same treatment group showed 5.4±2% cardiac injury (Fig. 2 B and C).

**Figure 2:**
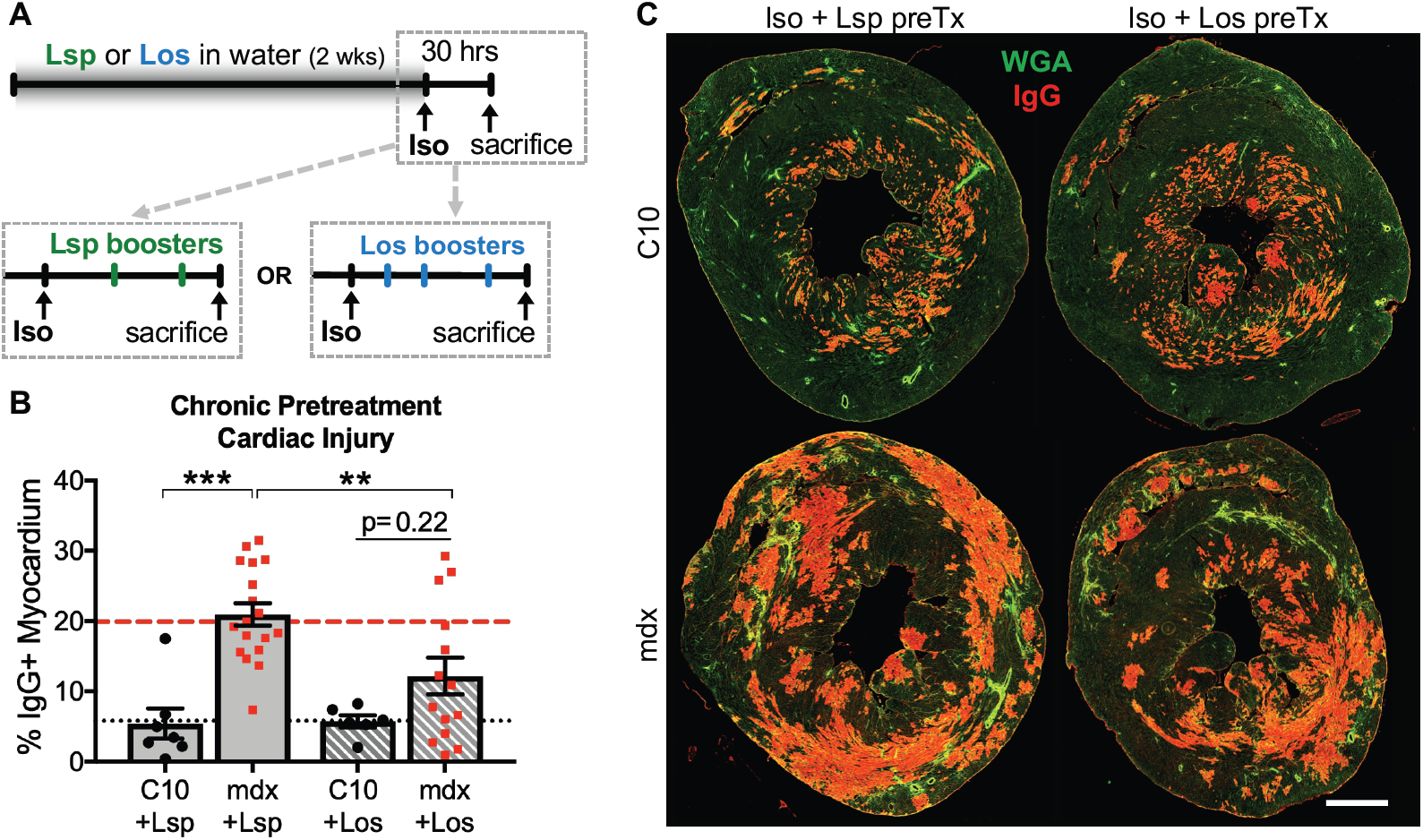
Pretreatment with losartan, but not lisinopril, protects *mdx* hearts from acute myocyte injury. **(A)** To investigate and compare potential chronic benefits of lisinopril (Lsp) and losartan (Los), each drug was given as a pretreatment in drinking water for 2 weeks before resuming the injection-based 30-hour protocol. **(B)** Losartan pretreatment significantly lowered Iso-induced IgG+ injury in *mdx* hearts, but did not alter the injury outcome in C10 hearts. Lisinopril pretreatment failed to show an effect in either group. Red dashed line indicates injury level of Iso-only *mdx* hearts, and black dotted line indicates injury in Iso-only C10 hearts, as shown in Fig. 1. (***p<0.001, **p=0.004; C10 n=6–7 per group; *mdx* n=14–18 per group). **(C)** Representative images of data in panel B. Whole heart transverse sections are shown with WGA (green) marking the tissue area and IgG (red) indicating areas of myocardial injury. Scale bar = 1 mm.

### 3.3. Patient records show that few patients with DMD receive ARB-based therapies

According to electronic records from patients with DMD seen in the years between 2009-2019 at the University of Minnesota, the majority of patients whose records mention an anti-angiotensin drug are receiving ACE inhibitors. Among the records that indicated the use of one of these drug classes, 72% of patients were on ACEIs while only 28% were taking ARB drugs (Fig. 3). The ACE inhibitor group included several patients that had begun the treatment before the age of 10, but all patients in the ARB treatment group were above the age of 10. Interestingly, only 38% (18/47) of patients in the ACEI-treated group but 72% (13/18) of the patients receiving ARBs were also being treated with beta blockers. This may be related to the fact that patients receiving ARBs were on average significantly older than those receiving ACEI, and thus more likely to be in more advanced stages of cardiac dysfunction (Fig. 3). Out of 95 patient records listing a diagnosis of DMD, 30 lacked any mention of ACE inhibitors or ARBs, and were thus excluded from the analysis.

**Figure 3:**
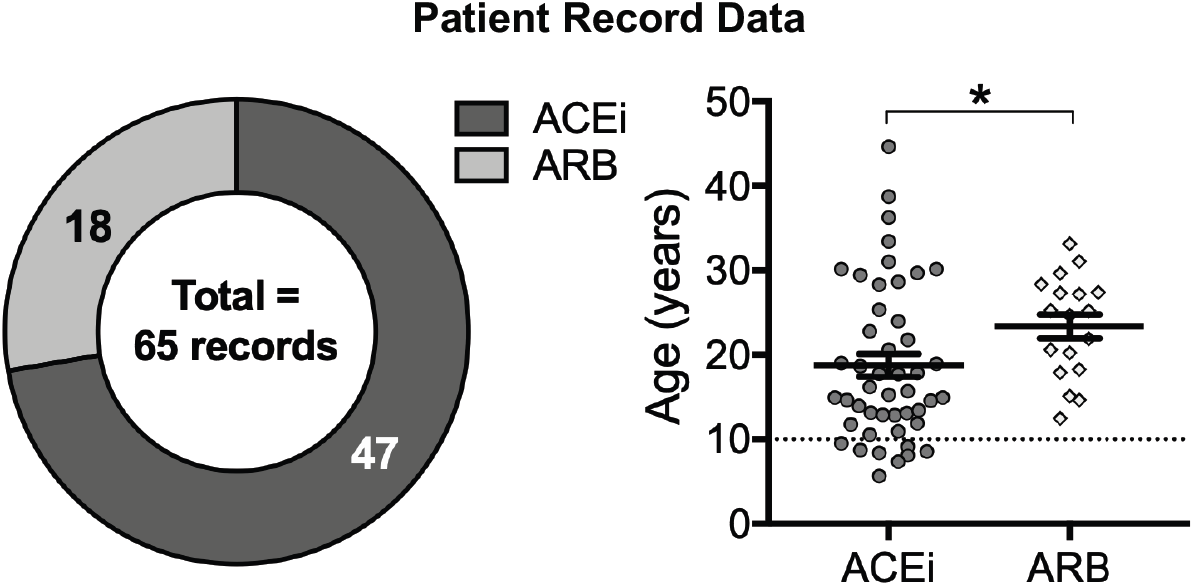
Patient records show that fewer patients with DMD receive treatment with ARBs than ACE inhibitors. *Left:* Of 65 records belonging to patients diagnosed with DMD that mentioned an anti-angiotensin drug, 72% (47/65) indicated the use of an ACEi, and 28% (18/65) indicated the use of an ARB. *Right:* The dotted line at 10 years of age corresponds to the 2015 guideline to initiate treatment with an ACEi or ARB at or before 10 years of age. Only the ACEi group includes patients who are younger than 10 years of age, and the group of patients receiving ARBs is older on average than the group receiving ACEi (*p=0.02).

## 4. Discussion

Cardiomyopathy is a major feature of DMD that afflicts essentially all patients over 18 years of age, becoming a leading cause of death in this era of life-prolonging symptomatic respiratory therapies [3,25,26]. The earliest signs of cardiac pathology appear in childhood, with transient elevations in serum levels of cardiac troponins and early cardiac fibrosis detected in patients as young as 6 years of age [4–6,27,28]. These observations indicate that the destruction of healthy myocardium is ongoing well before the onset of global cardiac dysfunction manifesting in reduced ejection fraction [4,28]. Therefore, clinical approaches that preserve the myocardium by preventing this early degeneration are critically important for the successful management of dystrophic cardiomyopathy. This evidence of early cardiac degeneration has resulted in the updating of earlier recommendations, that cardiac treatment be initiated in response to detectable deficits in function, with the current guidelines that all DMD patients without contraindications should start on ACE inhibitors or ARBs by 10 years of age or earlier [7,29]. Currently, most DMD patients receiving an anti-angiotensin therapy are on ACEIs, which are considered to be a first-line therapy for heart failure in general and are named first by the guidelines for managing dystrophic cardiomyopathy [7,8,30]. ARBs are often mentioned as an alternative to ACEIs or the next step for those who are poorly tolerant of ACEI side effects, although clinical trials have not shown any differences in efficacy between the two drugs in DMD [16,17].

This study and our previous work demonstrated that losartan treatment significantly lowered myocardial injury following Iso-induced cardiac stress in the *mdx* mouse heart. This injury-preventing effect is only evident in the dystrophic hearts, without significant reduction in wild type cardiac damage after losartan treatment. This observation suggests that an angiotensin-related mechanism that is uniquely exaggerated in the dystrophic process is driving the accumulation of myocardial injury in the dystrophic heart. This finding raised many questions regarding the potential ability of ACE inhibitors to deliver similar protection, prompting the work presented here. Surprisingly, we found a lack of benefit associated with lisinopril treatment before the onset of Iso-induced myocardial injury, regardless of the treatment being initiated 1 hour or 2 weeks before injury. These results indicate that ARBs have a unique protective action in the dystrophic heart that is not recapitulated by ACEI treatment.

Since ACE inhibitors and ARBs exert their respective actions at different points within the renin-angiotensin system, it may be expected that these drug classes would have similar beneficial effects. Indisputably, both drugs do provide significant benefits in the dystrophic heart, as indicated by multiple clinicals trials supporting their efficacy. However, the path from angiotensin I to AngII and AT_1_R activation is complex and multifaceted, opening the door for some divergence in the effects and benefits of these two drug classes that could explain the differences reported here. One possibility is that some combination of ACE-independent production of AngII by other peptidases and stretch-dependent activation of AT_1_R by the mechanical forces induced by increased cardiac workload results in substantial AT_1_R signaling even in the presence of ACEIs [31–35]. If the maximum benefit of anti-angiotensin therapies to cardiomyocytes lies in blocking AT_1_R activation, then the indirect inhibition provided by ACEIs may be insufficient to protect cardiomyocytes from damage in light of these alternative activating mechanisms.

Another possible explanation is that ARB and ACEI treatments trigger distinct homeostatic responses that result in different secondary effects on the production and metabolism of angiotensin peptides. One unique secondary effect that may contribute to the distinctive ability of ARBs to prevent cardiomyocyte injury may lie in enhanced activation of angiotensin II type 2 receptor (AT_2_R) as a compensatory consequence of AT_1_R antagonism. When AT_1_R is blocked, homeostatic mechanisms feed back to increase circulating levels of AngII, and all of this displaced AngII can serve as a ligand for AT_2_R activation, which has been shown to mediate cardioprotective effects of angiotensin peptides [35,36]. Conversely, the distinct secondary effects induced by ACEIs include increases in angiotensin (1-7) and bradykinin, both of which have documented cardioprotective actions that may play a role in the demonstrated clinical efficacy of ACEIs. Angiotensin (1-7) may be produced at higher rates by non-ACE peptidases secondary to the accumulation of angiotensin I that follows ACEI treatment, and may also be removed at a slower rate because its degradation is normally mediated by ACE, resulting in its accumulation with ACE inhibition [35,37]. Angiotensin (1-7) is the endogenous ligand for the Mas receptor, and like AT_2_R, Mas receptor activation causes a variety of protective effects that oppose the actions of AT_1_R, including reducing inflammation, reducing oxidative stress, and increasing vasodilation [38–40]. Bradykinin, the ligand for BK receptors, is another substrate for degradation by ACE that accumulates during ACE inhibition and may bear protective effects. However, while bradykinin is indeed linked to beneficial effects on inflammation, NO signaling, and pain, it is also a major contributor to the poorer tolerance of ACEIs, triggering a cough that prompts many patients to discontinue ACEI use [41]. Together, the joint effects of these protective mechanisms may drive the documented ACEI benefits of preventing remodeling and preserving function in the surviving myocardium. However, the inability of ACEIs to prevent myocardial damage in the Iso-induced injury model used here suggests that its protective mechanisms may be insufficient to overcome the acute effects of ACE-independent AT_1_R activation.

The work presented here has some important limitations that should be addressed in future studies. First, the molecular effects of ACEI and ARB that are described above participate in overlapping physiological pathways, thus it is not possible to estimate the magnitude of their respective contributions to the overall protection derived from the use of these drugs in a whole animal model. Further work should make use of more selectively targeted agonists and antagonists to investigate the distinct mechanisms that contribute to ACEI and ARB cardiac effects, including AT_1_R modulation, AT_2_R activation, and Ang 1-7 signaling, to better understand the protective pathways in the dystrophic heart. Furthermore, the studies described here have used the mildly-affected *mdx* mouse, so additional work will be needed to determine if these protective actions of ARBs are present in more severe or earlier-onset disease models. It is possible that a different approach to evaluating cardiac or skeletal muscle vulnerability in a dystrophic animal model would reveal a different relationship between the benefits of ACEI and ARB treatments.

Despite these limitations, the existence of the novel and specific protective influence of ARBs that we have described may carry major implications for clinical management of dystrophic cardiomyopathy. Not only does the injury-limiting capacity of ARBs support earlier initiation of anti-angiotensin therapies to precede the beginning of cardiac deterioration, but it also calls for a more thorough comparison of potential benefits between ARB and ACEI treatments. To date, clinical trials comparing ACEIs and ARBs have been performed in DMD patients old enough to have established cardiac injury, and records matching a DMD diagnosis from the University of Minnesota show a trend toward older patients in the ARB treatment group than the ACEi group. Based on these limitations, the lack of evidence supporting a significant benefit of ARBs over ACEIs may reflect the possibility that the additional ARB benefits described here are less evident in hearts with significant pre-existing myocardial damage. Additional clinical trials that enroll patients with DMD at a sufficiently young age to benefit from the potential protective actions of ARBs will be necessary to compare the therapeutic potential of ARBs and ACEIs in these fragile hearts. Data from our center show that the small proportion of patients currently receiving ARB treatment may already be too old to benefit maximally from early myocardial preservation by ARBs, so any comparisons based on data from patients who started their ARB treatment after the onset of cardiomyopathic symptoms should be made very cautiously.

In summary, we have presented data demonstrating that ARB treatment has a unique capacity to save cardiomyocytes from injury rather than just mitigating remodeling after their loss. These results support a shift in clinical practice toward using ARBs as a first-line therapy before 10 years of age for any child diagnosed with DMD, considering the certainty that virtually all of them will develop cardiomyopathy by their late teens. ARBs are safe, relatively well-tolerated, and readily available drugs that may hold the key to significant improvements in prognosis and quality of life for these patients, only requiring some adjustments to their clinical application.

## 5. Acknowledgments

The authors would like to thank Dr. Peter Karachunski and Dr. James Ervasti for helpful comments regarding the manuscript. This work was supported by the National Institutes of Health (R01 HL114832 and K08HL102066 to DT, F31HL139093 to TAM) and the Muscular Dystrophy Association (Grant 351960 to DT). The patient data collection was supported by National Institutes of Health’s National Center for Advancing Translational Sciences, grant UL1TR002494.

